# The oncoprotein DEK affects the outcome of PARP1/2 inhibition during replication stress

**DOI:** 10.1101/555003

**Authors:** Magdalena Ganz, Christopher Vogel, Christina Czada, Vera Jörke, Rebecca Kleiner, Agnieszka Pierzynska-Mach, Francesca Cella Zanacchi, Alberto Diaspro, Ferdinand Kappes, Alexander Bürkle, Elisa Ferrando-May

**Affiliations:** Department of Biology, Bioimaging Center, University of Konstanz, Konstanz, Germany; Nanoscopy and NIC@IIT, Istituto Italiano di Tecnologia, Genoa, Italy; Department of Experimental Oncology, European Institute of Oncology, Milan, Italy; Biophysics Institute (IBF), National Research Council (CNR), Genoa, Italy; DIFILAB, Department of Physics, University of Genoa, Genoa, Italy; Xi’an Jiaotong-Liverpool University, Dushu Lake Higher Education Town, Suzhou, China; Department of Biology, Molecular Toxicology Group, University of Konstanz, Konstanz, Germany

## Abstract

DNA replication stress is a major source of genomic instability and is closely linked to tumor formation and progression. Poly(ADP-ribose)polymerases1/2 (PARP1/2) enzymes are activated in response to replication stress resulting in poly(ADP-ribose) (PAR) synthesis. PARylation plays an important role in the remodelling and repair of impaired replication forks, providing a rationale for targeting highly replicative cancer cells with PARP1/2 inhibitors. The human oncoprotein DEK is a unique, non-histone chromatin architectural protein whose deregulated expression is associated with the development of a wide variety of human cancers. Recently, we showed that DEK is a high-affinity target of PARylation and that it promotes the progression of impaired replication forks. Here, we investigated a potential functional link between PAR and DEK in the context of replication stress. Under conditions of mild replication stress induced either by topoisomerase1 inhibition with camptothecin or nucleotide depletion by hydroxyurea, we found that the effect of acute PARP1/2 inhibition on replication fork progression is dependent on DEK expression. Reducing DEK protein levels also overcomes the restart impairment of stalled forks provoked by blocking PARylation. Non-covalent DEK-PAR interaction via the central PAR-binding domain of DEK is crucial for counteracting PARP1/2 inhibition as shown for the formation of RPA positive foci in hydroxyurea treated cells. Finally, we show by iPOND and super resolved microscopy that DEK is not directly associated with the replisome since it binds to DNA at the stage of chromatin formation. Our report sheds new light on the still enigmatic molecular functions of DEK and suggests that DEK expression levels may influence the sensitivity of cancer cells to PARP1/2 inhibitors.

## INTRODUCTION

Poly(ADP-ribosyl)ation (PARylation) is an abundant protein posttranslational modification regulating numerous cellular functions among which the maintenance of genomic stability plays a prominent role [1]. The enzyme responsible for 85-90% of the cellular PAR synthesis activity is PARP1, with PARP2 accounting for the remainder [2]. PAR can be covalently linked to and/or interact non-covalently with target proteins. PARylation is highly dynamic and can be very transient in nature due to the activity of the de-modifying enzyme, the PAR glycohydrolase or PARG [3]. Inhibition of PARylation by small molecule compounds is a recently approved strategy for the treatment of ovarian cancer [4]. The rationale for the use of PARP1/2 inhibitors in chemotherapy is based on their synthetic lethal interaction with DNA damaging agents in cells which are deficient for recombinational DNA repair through mutations in BRCA1/2 [5, 6]. In these cells, inhibition of PARylation abrogates base excision repair thereby turning endogenous single strand breaks (SSBs) in highly toxic, non-repairable double strand breaks (DSBs). In addition, PARP1/2 inhibitors possess DNA trapping activity which causes DSBs on its own due to the collision of PARP-DNA complexes with the DNA replication and transcription machineries [7]. Impaired DNA replication has recently come into the focus as a further source of DNA lesions which can become lethal to cells treated with PARP1/2 inhibitors. If not removed timely, replication blocks lead to fork collapse leaving behind single ended DNA strand breaks as well as SSBs which require PARylation for their prompt repair. PARP1/2 was also shown to be directly involved in replication fork stabilization and protection. Thus, PARP is required for the restart of collapsed forks after prolonged exposure to hydroxyurea (HU) [8], protects transiently stalled forks from premature and extensive resection [9] and regulates fork reversal induced e.g. by low doses of camptothecin (CPT). More precisely, PARylation prevents RecQ helicase from resolving regressed forks prematurely, thus avoiding fork run off across DNA lesions and DSB generation [10, 11]. Finally, PARP1/2 was shown to play an important role also during unperturbed DNA replication. Using pharmacological PARG inhibition to stabilize and detect basal PAR levels, the polymer was shown to be required for sensing and repairing a sub-set of unligated Okazaki fragments thus providing a back-up pathway for the completion of lagging strand DNA synthesis [12].

DEK is a non-histone chromatin protein which is ubiquitously present in higher eukaryotes [13]. Its binding to DNA is regulated by abundant post-translational modifications, including phosphorylation [14, 15], acetylation [16, 17], and PARylation [18–20]. Covalent PARylation of DEK is efficiently triggered by DNA damage leading to the loss of its DNA binding and folding activities [20]. The DEK amino acid sequence bears three PAR-binding motifs which mediate non-covalent PAR interaction *in vitro*, thereby moderately reducing DNA binding but incrementing DEK multimerization [18]. A continuously increasing number of studies link DEK overexpression to cancer development, pinpointing DEK as a “bona fide” oncogene [21]. DEK is considered a potential therapeutic target and a biomarker for breast and ovarian cancer [22–24], retinoblastoma [25], colorectal [26] and bladder cancer [27] as well as for melanoma progression [28, 29]. DEK has a pleiotropic mode of action and can influence diverse regulatory circuits in the cell, a notion supported also by the recent elucidation of its interactome [30]. Downregulation of DEK expression increases the susceptibility to DNA damage [20, 31], attenuates apoptosis [32] and senescence [21], and affects proliferation and chemoresistance [23, 33, 34]. On the mechanistic level, DEK is known to have DNA folding activity, principally via its ability to introduce positive supercoils [35–37]. Thus, DEK has been involved in splicing [38], transcriptional activation and repression, heterochromatin stability [39], DNA repair [40, 41] and DNA replication [42, 43]. Concerning the latter, we recently proposed that DEK acts as a tumour promoter by protecting cells from the deleterious consequences of DNA replication stress. In particular, we showed that DEK facilitates replication fork progression under stress, and counteracts DNA damage arising from impeded replication as well as its transmission to daughter cells [43]. In this study, we set out to examine a potential functional link between DEK and PARP1/2 in the context of DNA replication stress. Our data reveal that for mildly stressed replication forks, the consequences of PARP1/2 inhibition depend on DEK expression.

## MATERIAL AND METHODS

### Cell culture

U2-OS osteosarcoma cells were cultured in McCoy’s 5a modified medium (Thermo Fisher Scientific) supplemented with 10 % fetal bovine serum (FCS; Capricorn Scientific and PAA Laboratories), 100 U/ml penicillin and 100 μg/ml streptomycin (both Thermo Fisher Scientific). U2-OS control and shDEK cells [43] were additionally supplemented with 2 μg/ml puromycin (Merck). Puromycin was omitted 36 h prior to experiments. U2-OS shDEK cells stably express an shRNA targeting the human DEK transcript, resulting in a permanent reduction of DEK protein levels of around 90 % [43]. U2-OS wildtype cells were a kind gift of G. Marra, University of Zurich, Switzerland. To generate U2-OS GFP-DEK cells, the eGFP sequence has been inserted at the 5`end of the endogenous DEK coding sequence in wildtype cells via TALEN-mediated genome editing (Vogel et al., in preparation). HeLa S3 cervical adenocarcinoma cells were cultured in DMEM medium (Thermo Fisher Scientific) supplemented with 10 % fetal bovine serum, 100 U/ml penicillin, 100 μg/ml streptomycin and 6 mM L-glutamine (Thermo Fisher Scientific). BJ-5ta foreskin fibroblasts were cultured in a 4:1 mixture of DMEM medium and Medium 199 medium (Thermo Fisher Scientific) supplemented with 10 % fetal bovine serum, 4 mM L-glutamine and 10 μg/ml hygromycin B (Merck).

For induction of replication stress, cells were treated with hydroxyurea (HU; Merck) or camptothecin (CPT; Merck) as indicated. PARP1/2 activity was inhibited with ABT-888 or AZD-2281 (both Selleckchem) as indicated.

### Isolation of Proteins on Nascent DNA (iPOND)

iPOND was performed as described by Sirbu et al. [44], with minor modifications. At least 1×10^8^ HeLa S3 cells per sample were pulsed with 10 μM EdU (Thermo Fisher Scientific) for the indicated times and either incubated with 10 μM thymidine (Merck) for 0 - 30 min before fixation (chase experiments) or fixated immediately (pulse experiments). For replication stress experiments thymidine containing medium was supplemented with 2 mM HU and/or 1 μM ABT-888. Click reaction to label EdU-containing DNA was performed using biotin-PEG3-azide (Jena Bioscience) for 90 min and cells were sonicated in a Bioruptor sonicator (Diogenode) to solubilize chromatin fragments. Biotin-linked fragments were precipitated overnight at 4 °C using streptavidin-coupled magnetic beads (0.8 μm, Solulink). Chromatin bound proteins (“Capture”) were subjected to Western blot analysis using the following antibodies: polyclonal rabbit α-DEK K-877 (1:20,000; [20]), monoclonal mouse α-PCNA (1:9,000; PC10, Cell Signaling Technology), polyclonal rabbit α-H3 (1:150,000; ab1791, Abcam).

### DNA fiber assay

For the determination of tract length ratios, U2-OS shDEK and control cells were labelled with 60 μM CldU (Merck) for 20 min and subsequently treated with 250 μM IdU (Merck) for 20 min in the presence or absence of 25 mM CPT and/or 1 μM ABT-888 as indicated. For the analysis of replication fork restart, cells were labelled with 60 μM CldU for 20 min and subsequently treated with 4 mM HU and 60 μM CldU for 4 h. After washing, cells were labelled with 250 μM IdU for 20 min in the presence or absence of 1 μM ABT-888.

DNA fiber spreads were prepared as described by Merrick et al. [45] with modifications: After trypsination and resuspension in ice-cold PBS, labelled and unlabelled cells were mixed in a 1:5 ratio. 12.5 μl of the mixture were diluted with 7.5 μl lysis buffer (200 mM Tris-HCl pH 7.4, 50 mM EDTA, 0.5 % SDS) on a glass slide. After 9 min the slides were tilted at 30-45° and the resulting DNA spreads were air-dried and fixed overnight in a methanol:acetic acid mixture (3:1) at 4 °C. Following denaturing and blocking with 2 % BSA in 0.1 % Tween 20, the slides were incubated for 2.5 h with rat α-BrdU (1:200; BU1/75 (ICR1), Abcam; detects CldU) and mouse α-BrdU (1:200; B44 from BD Biosciences; detects IdU) antibodies. Fibers were treated with goat α-mouse AlexaFluor-488 and goat α-rat AlexaFluor-546 (both Thermo Fisher Scientific) secondary antibodies for 1h at RT, allowed to air-dry and mounted in ProLong Gold Antifade (Thermo Fisher Scientific). Widefield microscopy was performed with a Zeiss Axio Observer Z1 equipped with a Plan Apochromat 63x/1.40 oil DIC objective lens. Data were evaluated using Fiji v1.49u (National Institutes of Health, MD [46]) and the fiber tool of the BIC macro tool box [43].

### Selection of DEK-GFP expressing cells

U2-OS shDEK cells were transfected with plasmids encoding DEK WT-GFP or DEK PBD2-Mut2-GFP. After 24 h, cells were sorted using a FACSAria Illu (BD Biosciences). Low DEK GFP expressing cells were collected in McCoy’s 5a modified medium supplemented with 20 % FCS. To determine expression levels of endogenous and ectopic DEK, total proteins were extracted with SDS lysis buffer. Cleared lysates were subjected to Western blotting with the following antibodies: polyclonal rabbit α-DEK K-877 (1:20,000; [20]), polyclonal rabbit α-PCNA (1:5,000; ab18197, Abcam).

### Immunofluorescence

For immunofluorescence detection of Rad51, U2-OS cells were preextracted using CSK-buffer (10 mM Hepes-KOH pH 7.4, 300 mM sucrose, 100 mM NaCl, 3 mM MgCl2, 0.5 % Triton X-100, 10 mM NaF, 1 mM NaVO3, 11.5 mM Na-molybdat) for 5 min on ice after treatment and fixed using 4 % PFA/PBS supplemented with 10 mM NaF and 1 mM NaVO3 (20 min, RT). For immunofluorescence detection of 53BP1, γH2AX and RPA70, cells were fixed with 4 % PFA/PBS without preextraction. After permeabilization, cells were incubated with primary antibodies diluted in 1 % BSA/PBS (Rad51, 53BP1, γH2AX) or 10 % FBS/0.05 % Na-azide/culture medium (RPA70) overnight at 4 °C. The following primary antibodies were used: polyclonal rabbit α-53BP1 (1:200; H-300, Santa Cruz), monoclonal rabbit α-RPA70 (1:1000; ab79398, Abcam), monoclonal mouse α-γH2AX (1:500; Ser139, clone JBW301, Santa Cruz), polyclonal rabbit α-Rad51 (1:100; H-92, Santa Cruz). After washing with PBS, cells were incubated with secondary antibodies diluted in 1 % BSA/PBS at RT for 1 h. The following secondary antibodies were used: goat α-mouse AlexaFluor-488, goat α-rabbit AlexaFluor-546 (both 1:400; both Thermo Fisher Scientific). For nuclear counterstaining, cells were incubated in 200 ng/μl Hoechst 33342/PBS (Merck). Coverslips were mounted on microscopy slides using Aqua Polymount (Polysciences). Replicating cells were visualized by labelling with 10 μM EdU 10 min prior to replication stress induction. Cells were fixed and immunostainings performed as described above. After incubation with secondary antibodies, EdU was detected using the Klick-it EdU Imaging Kit with AlexaFluor-488 or -647 azide (all Thermo Fisher Scientific) following manufacturer’s instructions. Nuclear counterstaining and mounting of coverslips was performed as described above.

### Confocal and superresolution microscopy

Confocal microscopy was performed with a Zeiss LSM 510 Meta and a Zeiss LSM 780 equipped with a Plan Neofluar 40x/1.30 oil or a Plan Apochromat 40x/1.40 oil objective lens, respectively. Image analysis was done with Fiji v1.49u [46] using the ImageJ BIC macro tool box [43]. For counting DNA damage foci, appropriate noise parameters for each channel were determined manually and applied to all samples within one experiment. For the determination of the number of cells positive for lesion markers, the lower threshold for the number of foci per nucleus was set such to include 95 % of untreated control cells. The threshold was applied to all samples within one experiment. Cells exceeding the threshold were classified as positive for the respective lesion marker.

To test the efficiency of PARP inhibitor ABT-888, cells were left untreated or pre-treated with 1 μM ABT-888 for increasing time points, followed by DNA damage induction using 800 μM H_2_O_2_ (Merck) for 10 min. Detection of PAR was achieved after fixation with methanol:acetic acid (3:1) using a monoclonal mouse α-PAR antibody (10H, 1:300 in PBSMT (5 % milk powder, 0.05 % Tween 20, PBS) at 4 °C overnight. Confocal microscopy was performed with a Zeiss LSM 510 Meta as described above. PAR nuclear intensities were analysed using Fiji v1.49u.

For superresolution imaging by 3D structured illumination (SI), U2-OS GFP-DEK cells were grown on high precision coverslips (# 1.5). Replication foci were labelled via incubation with 10 μM EdU for 10 min prior fixation. For immunofluorescence detection of PCNA, cells were fixed using 4 % PFA/PBS (10 min) and permeabilized in methanol (5 min, −20 °C). After blocking with 2 % BSA/PBS, cells were incubated with monoclonal mouse α-PCNA primary antibody (PC10, Cell Signaling Technology) diluted 1:2,400 in 10 % NGS/PBS at 4 °C overnight. After washing with 0.05 % Tween/PBS, cells were incubated with goat α-mouse AlexaFluor-568 secondary antibody diluted 1:400 in 10 % NGS/PBS (1 h, RT). After washing with 0.05 % Tween/PBS, cells were fixed again using 2 % PFA/PBS (10 min, RT). EdU was detected using the Klick-it EdU Imaging Kit with azide AlexaFluor-647. Coverslips were mounted on microscopy slides using Vectashield H-1000 (Vector Laboratories). Images were acquired at a DeltaVision OMX Blaze v4 (GE Healthcare) using an Olympus Plan Apochromat 60x/1.42 oil objective. A z-stack of at least 20 slices (0.125 μm step size) was acquired per image in SI mode. Reconstruction of SIM images and image registration of the channels was performed using softWoRx v6.5.2 (GE Healthcare). Pseudo-widefield images were generated with Fiji v1.51n and the SIMcheck plugin (v1.0 [47]).

Super resolved imaging by stochastic optical reconstruction microscopy (STORM) was carried out on U2-OS GFP-DEK cells cultured in McCoy’s 5a modified medium (Thermo Fisher Scientific) supplemented with 10 % fetal bovine serum (FCS; Capricorn Scientific and PAA Laboratories), 100 U/ml penicillin and 100 μg/ml streptomycin (both Thermo Fisher Scientific). For STORM imaging cells were plated on eight-well Lab-Tek coverglass chamber (Nunc), grown under standard conditions and fixed after 24 h. For STORM detection of DEK protein cells were fixed with methanol:ethanol (1:1) at −20 °C for 3 min, washed with PBS following blocking buffer for 2 h at RT (3 % BSA, 0.2% Triton X-100, PBS). After blocking and permeabilization, cells were incubated with primary antibodies diluted in blocking buffer, firstly for 2 h at RT and then overnight at 4 °C. The following primary antibodies were used: polyclonal chicken α-GFP (1:2000; ab13970, Abcam), polyclonal rabbit α-PCNA (1:50; HPA030522, Sigma-Aldrich). After six washing steps with blocking buffer for 5 min each, cells were incubated with secondary antibodies diluted in blocking buffer at RT for 1 h. Specific antibodies were used, namely: donkey α-chicken AlexaFluor405/AlexaFluor647 (1:50; IgG (703-005-155, Jackson ImmunoResearch) coupled to reporter and activator dyes – AlexaFluor405 #A30000, Invitrogen and AlexaFluor647 #A2006), and goat α-rabbit CF568 (1:1000; SAB4600084, Sigma-Aldrich).

Single-molecule localisation was performed using a Nikon N-STORM super-resolution microscope equipped with a 100x/1.40 oil-immersion objective lens and coupled to an Andor iXon DU-897E-CS0BV EMCCD camera (image pixel size 160 nm) with 30ms exposure time. To maintain the z-position a Nikon “perfect focus system” was used. The set-up included a 405 nm laser for activation (Coherent CUBE 405 nm; 100 mW) and a 647 nm readout laser (MPBC’s CW Visible Fiber Laser). Imaging was performed using TIRF illumination. 30.000 frames at 25 Hz frame rate were acquired. For widefield imaging together with STORM, a 561 nm laser (Coherent Sapphire OPSL 561 nm; 100 mW) was used. Dichroic mirrors and band-pass filters allowed for selection of emitted signals (ZET405/488/561/647, Chroma). For super-resolution measurements, STORM imaging buffer was used (prepared following Nikon’s STORM Protocol-Sample Preparation) containing GLOX solution as oxygen scavenging system (40 mg/ml Catalase, Sigma; 0.5 mg/ml glucose oxidase; 10 % glucose in PBS) and MEA 10 mM (Cysteamine MEA, Sigma-Aldrich, #30070-50G, in 360 mM Tris-HCl). Single molecule localization and super-resolution image reconstruction were performed using NIKON software (NIS elements) and a custom software (Insight3, custom software developed by B. Huang, University of California). Molecules are identified and x-y located by Gaussian fitting. The final image is reconstructed, after drift correction, by plotting each identified molecule as a Gaussian spot with a width corresponding to the achieved localization precision (9nm).

### Site-directed mutagenesis

Nucleobase mutations of the DEK primary sequence were introduced via a modified Quick ChangeTM site-directed mutagenesis protocol [48]. To generate the DEK-GFP template for the mutagenesis PCR, the DEK WT sequence was inserted into an eGFP reporter plasmid (peGFP-N1, Addgene 6085-1). PAR-binding domain 2 (bases 583 - 663) and the DEK shRNA target site (bases 1000 – 1020) were mutated using overlapping primer pairs containing the desired base changes. For the PAR-binding domain a total of 9 codons were mutated in four rounds of mutagenesis (primers PBD2-Part1-4, see also Fig 7 B). For the shRNA target site a total of 8 nucleobases were mutated in two rounds (primers shDEK-Part1-2), resulting in silent mutations and diminished binding of the DEK shRNA. Primer sequences are listed in S1 Table.

### Expression and purification of recombinant GST-tagged proteins from *E. coli*

The mutated DEK sequence (DEK PBD2-Mut2) was inserted into a GST expression plasmid. LB_amp_ medium was inoculated with *E.coli* BL21(DE3) pGEX 4T-1 harbouring plasmids encoding GST only, GST-DEK WT, or GST-DEK PBD2-Mut2. Protein expression was induced using 0.5 mM IPTG (Merck) for 1.5 h. Bacteria were harvested via centrifugation (1,600 × g, 15 min, 4 °C), the pellet was resuspended in resuspension buffer (20 mM Tris-HCl pH 8, 1 M NaCl, 0.5 mM EDTA, 1 mM DTT) and shock frozen in liquid nitrogen. Cells were sonicated on ice, 0.5 % NP-40 (Merck) was added and the lysate was centrifuged (18,000 × g, 30 min, 4 °C). The supernatant was incubated with 200 μl Glutathion-Sepharose 4B-beads (GE Healthcare) equilibrated in wash buffer I (20 mM Tris-HCl pH 8, 500 mM NaCl, 0.5 mM EDTA, 1 mM DTT, 0.1 % NP-40) for 2 hours at 4 °C. Beads were washed with wash buffers of decreasing NaCl concentrations (500, 300 and 20 mM NaCl). For elution of GST-tagged DEK, beads were incubated with 200 μl of elution buffer (200 mM Tris-HCl pH 8, 20 mM NaCl, 40 mM reduced glutathione, 10 % glycerol) for one hour at 4 °C. The GST-tagged protein containing supernatant was shock frozen in liquid nitrogen and stored at −80 °C. Protein concentrations were determined using the BCA Protein Assay Kit (Thermo Fisher Scientific) according to manufacturer’s instructions.

### Electrophoretic Mobility Shift Assay (EMSA)

Purified recombinant proteins (GST-DEK WT and GST-DEK PBD2-Mut2) were dialyzed in nE100 buffer (20 mM Hepes-KOH pH 7.6, 100 mM NaCl, 10 mM NaHSO_3_, 1 mM EDTA, supplemented with 1 μg/ml BSA) using Millipore filters (VSWP 0.025 μm; Merck) for 90 min at 4 °C. 175 ng of plasmid DNA were incubated with increasing amounts of recombinant DEK in a total volume of 30 μl nE100 buffer for one hour at 37 °C. Samples were subjected to electrophoresis on a 0.6 % agarose gel in TBE buffer (50 mM Tris base, 80 mM boric acid, 1 mM EDTA, pH 8). DNA-protein complexes were visualized with 0.5 μg/ml ethidium bromide solution using a fluorescence imager.

### *In-vitro* synthesis of PAR

PAR was synthesized and purified according to Fahrer et al. [49]. Briefly, 50 μg/ml ‘activator’ oligonucleotide GGAATTCC and 60 μg/ml of both recombinant Histone H1 und H2A were diluted in buffer containing 100 mM Tris-HCl pH 7.8, 10 mM MgCl_2_ and 1 mM DTT. To start the reaction, 1 mM NAD^+^ (Merck) and 150 nM recombinant PARP1 were supplemented. PAR synthesis was stopped after 15 min by adding ice-cold trichloroacetic acid (TCA) to a final concentration of 10 %. PAR was detached from histones and PARP1 itself using 0.5 M KOH/50 mM EDTA. After neutralization, DNA and proteins were digested using 110 μg/ml DNase and 220 μg/ml proteinase K (both Merck), respectively. PAR was purified by phenol-chloroform-isoamylalcohol extraction and ethanol precipitation. The concentration of the purified polymer was determined via absorbance at 258 nm.

### PAR overlay assay

60 pmol of custom synthesized PAR-binding domain peptides (for sequences see Fig 7 B and S 5 A Fig; biomers.net) or 25 pmol of recombinant GST-tagged proteins were transferred onto a nitrocellulose membrane using a slot blotting apparatus. The membrane was allowed to air-dry and incubated overnight in 5 pmol PAR/TBST (100 mM Tris-HCl, 150 mM NaCl, 0.5 % Tween 20) at 4 °C. Blots were blocked in 5 % milk powder/TBST and membrane-bound PAR was detected using monoclonal mouse α-PAR antibody (1:300; 10H). After washing, the membrane was incubated with secondary antibody goat α-mouse Ig/HRP (1:2,000; Agilent). PAR was detected using a chemiluminescence imager. To verify that equal amounts of proteins or peptides were blotted onto membranes, the same protein solutions as used for the PAR overlay were slot blotted onto a nitrocellulose membrane and allowed to air-dry. Samples were fixed in 7 % acetic acid/10 % methanol for 15 min at RT. After fixation, proteins or peptides were stained using Sypro Ruby Protein Blot Stain (Thermo Fisher Scientific) for 15 min and visualized using a Gel Doc XR system (Bio-Rad).

### Statistical analysis

Statistical tests were performed using GraphPad Prism 5.02 and applied as indicated in the figure legends. \**p* ≤ 0.05, \*\**p* ≤ 0.01, \*\*\**p* ≤ 0.001.

## RESULTS

### The effect of short term PARP1 inhibition on mildly challenged replication forks is reverted in DEK knockdown cells

To investigate whether PARylation regulates the impact of DEK on the replication stress response, we set out from our previous observation that downregulation of DEK expression aggravates replication fork slowing induced by low concentrations of CPT. Inhibition of topoisomerase1 by CPT stabilizes Topo1-cleavable complexes (Top1ccs), thus causing torsional stress ahead of the replication fork. As a result, fork progression is impaired, eventually leading to replisome disassembly and DNA strand breaks. At very low doses (25 nM), CPT was shown to slow down, but not arrest, fork progression and trigger fork reversal in a PARP1/2-dependent manner [10, 11].

Firstly, we examined the effect of DEK and PARP1/2 activity on CPT-induced replication fork progression using DNA fiber assays (Fig 1). Cells bearing a stable, lentiviral mediated knockdown of DEK expression (shDEK cells) and the respective control cells [43] were treated with 25 nM CPT in the presence and absence of ABT-888. Both drugs were added simultaneously with the IdU-containing medium (Fig 1 A). In line with our previous results, knockdown of DEK expression slowed down replication fork progression *per se* as indicated by a highly significant reduction of the IdU tract lengths in untreated shDEK cells as compared to controls (Fig 1 C and [43]). Inhibition of PARP1/2 activity with ABT-888 had no significant effect on replication fork speed in both control and shDEK cells, measured as the ratio of the IdU-labelled tracts (green) vs the CldU-labelled tracts (red) (Fig 1 D, compare boxes 1 and 2). Notably, in all our PARP1/2 inhibition experiments, we took care of minimizing DNA damage due to trapping of the enzyme on DNA [7]. Therefore, we used ABT-888 as an inhibitor with reportedly low trapping activity and limited the exposure to the duration of the IdU-pulse. PARP1/2 activity is effectively inhibited under these conditions (S1 A, B Fig) but does not trigger a DNA damage response, as indicated by the absence of 53BP1 foci formation (S1 C, D Fig; see also [50, 51]).

**Fig 1.**
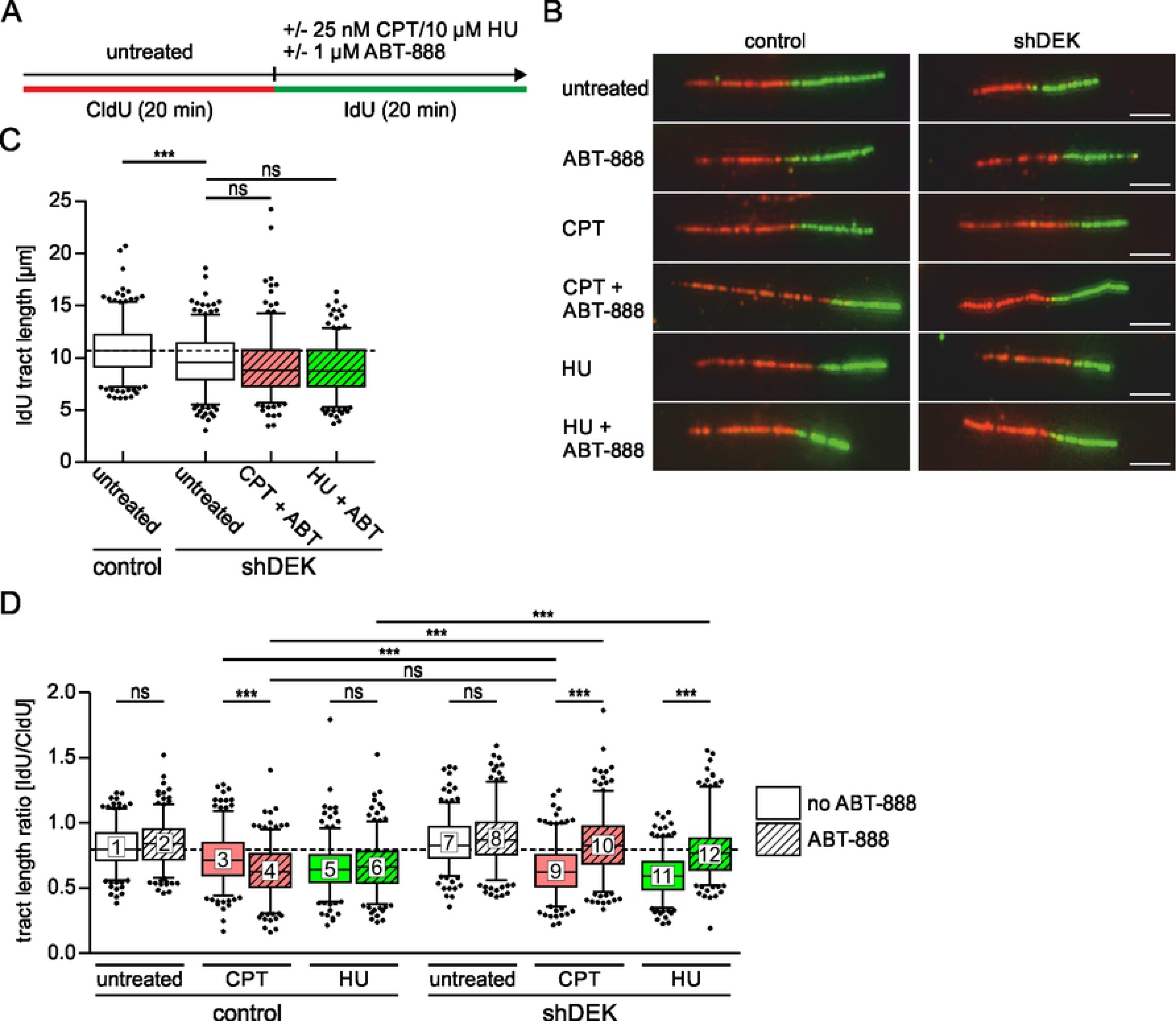
Combined inactivation of DEK and PARP1/2 prevents fork slowing by low doses of CPT and HU. (A) Scheme of the DNA fiber assay. U2-OS control and shDEK cells were pulse-labelled with CldU for 20 min, followed by incubation with IdU for 20 min in the presence or absence of replication stress inducers (25 nM CPT or 10 μm HU) and 1 μM ABT-888. (B) Representative microscopic images of DNA fibers after spreading. Thymidine analogues were visualized via indirect immunofluorescence. CldU-labelled tracts were visualized in the red channel, IdU-labelled tracts in the green channel. Scale bar: 5 μm. (C-D) Quantification of CldU and IdU tract lengths of at least 250 fibers per experimental condition. The experiment was performed in triplicates. The bands inside the boxes display the median, whiskers indicate the 5th to 95th percentile and black dots mark outliers. t-test: ns: not significant, *** p≤0.001. ABT-888 treated cells: hatched bars. (C) Lengths of IdU-labelled tracts. (D) IdU/CldU tract length ratios.

Treatment with low doses of CPT reduced fork progression, as expected, in control cells and, to a greater extent, in shDEK cells (Fig 1 D, compare boxes 3 and 9). When exposure to CPT occurred in the presence of ABT-888, the two cells lines, however, showed opposite responses: in control cells, the replication fork was further retarded as compared to treatment with CPT only, while in shDEK cells, fork speed recovered to the basal level measured in the absence of any perturbation (Fig 1 D, compare boxes 4 and 10). Further, in the presence of CPT, the extent of fork slowing obtained by inhibiting PARP1/2 in control cells equalled that resulting from downregulation of DEK expression (Fig 1 D, compare boxes 4 and 9), which is suggestive of DEK and PARP1/2 acting in the same regulatory pathway.

We sought to validate these observations in a different replication stress model and performed fiber assays using low doses of hydroxyurea (HU). At 10 μm, HU slows down but does not stall the replication fork as observed at higher concentrations (e.g. 2mM, compare Fig 1 D with S2 Fig). Again, additional PARP1/2 inhibition positively impacted on fork progression in shDEK cells, but not in control cells (Fig 1 D, compare boxes 6 and 12). Interestingly, this modulatory effect was detectable only under mild replication stress conditions. At a concentration of HU of 2 mM, combined exposure to ABT-888 did not alter fork speed, although in general, replication forks of shDEK cells were significantly more sensitive to HU-mediated stalling than those of control cells (S2 Fig). Finally, the fork acceleration observed in mildly stressed, PARylation inhibited shDEK cells was not sufficient to compensate for the fork impairment caused by DEK downregulation itself, because the IdU tract length in stressed and PARP1/2-inhibited shDEK cells remained shorter than in control untreated cells (Fig 1 C).

As replication stress is a source of DNA damage, we evaluated whether DEK downregulation would also affect the formation of DNA strand breaks caused by exposure to replication inhibitors and ABT-888. We assessed replication-associated DSBs by counting γH2AX/53BP1 double-positive foci in EdU-positive S-phase shDEK and control cells. In the case of CPT, DNA strand breaks are known to arise when the transcription and/or replication machineries collide with unrepaired Top1ccs. At 25 nM CPT, the overall response was very moderate as expected at this subtoxic dose [11]. Only 7 foci were observed on average in control cells (Fig 2 A, B). This low number was nevertheless significantly higher in shDEK cells, in line with our previous data showing that DEK downregulation sensitizes cells to CPT treatment [20]. The combination of CPT and PARP1/2 inhibition led to an increase in DSBs in control cells, while shDEK cells showed a significant reduction, resembling the pattern observed with our fiber assays. These data suggest that the restoration of fork speed observed in CPT-treated, DEK-depleted cells when PARP1/2 is inhibited is not a manifestation of fork run off. Most likely, this effect reflects the ability of shDEK cells to either withstand the action of CPT or better cope with its consequences, if PARP1/2 activity is blocked. Unfortunately, this assumption could not be confirmed in the HU-model of replication stress, since exposure to 10 μM HU did not result in a measurable DNA damage response in our experimental setting (Fig 2 A, B).

**Fig 2.**
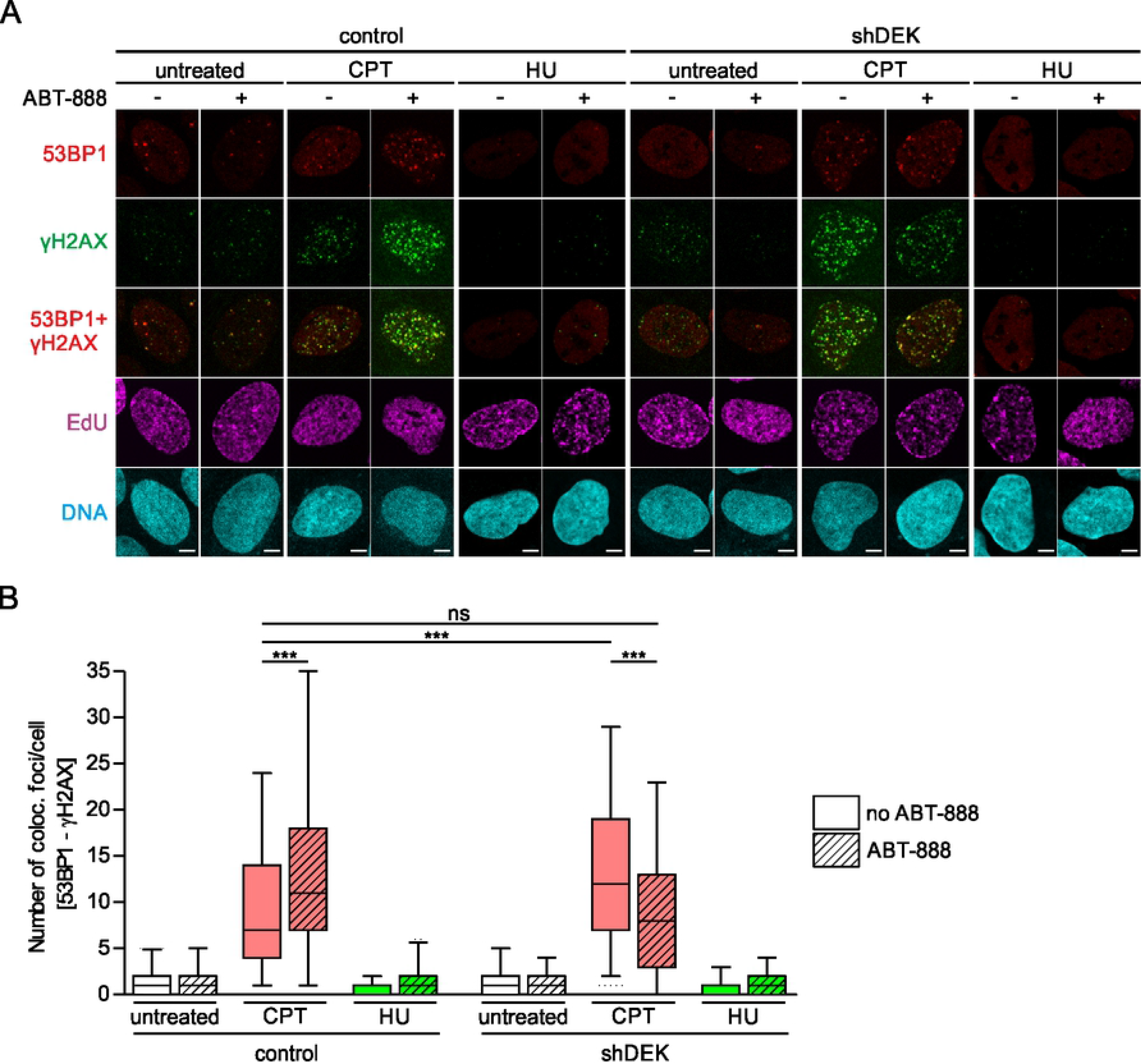
Combined inactivation of DEK and PARP1/2 counteracts DNA damage induced by low doses of CPT. U2-OS control and shDEK cells were pulse-labelled with EdU, then either left untreated or treated with replication stress inducers (25 nM CPT or 10 μm HU) for one hour, in the presence or absence of 1 μM ABT-888. 53BP1 (red) and γH2AX (green) foci formation was visualized via indirect immunofluorescence analysis, EdU (magenta) using click chemistry. DNA was counterstained with Hoechst 33342 (cyan). (A) Representative confocal images. Scale bar: 5 μm. (B) Quantification of 53BP1/γH2AX colocalization in S-phase cells. Foci were counted and colocalization determined using the foci counter of the BIC macro tool box. At least 118 cells per experimental condition were evaluated. The experiment was performed in triplicates. The bands inside the boxes display the median, whiskers indicate the 5th to 95th percentile and outliers are omitted for clarity. t-test: *** p≤0.001.

Altogether, these data show that the outcome of acute PARP1/2 inhibition on challenged replication forks is dependent on DEK expression levels and suggest the existence of a regulatory network involving DEK and PARP1/2 that modulates fork speed. This interaction is only detectable under mild stress levels, when the fork is still processive.

### DEK is not part of the replisome but binds to newly replicated DNA as it matures to chromatin

The marked effects of DEK expression on fork progression let us explore whether DEK directly associates to replication forks. To this end, we performed iPOND (isolation of proteins on nascent DNA) assays [44]. Firstly, we treated HeLa S3 cells with EdU for increasing time periods (2.5 min – 30 min) to label newly synthesized DNA and subsequently monitored the occurrence of DEK in the pool of enriched proteins by Western Blot (Fig 3 A-C). We used PCNA to monitor the active replisome while histone H3 served as marker for maturing chromatin. PCNA was detected at early time points (5 min and 10 min) representing nascent DNA. DEK lagged behind and appeared after an EdU pulse of 15 min duration, concomitantly with H3. We corroborated this result with an iPOND pulse-chase experiment, in which we sought to observe the dynamics of DEK binding to nascent chromatin (Fig 3 D-F). EdU was applied for 10 min to pulse-label nascent DNA and then replaced with thymidine for increasing time periods before the isolation of proteins crosslinked to DNA. Proteins binding directly and exclusively at the replication fork diminish in the enriched protein fraction as the EdU labelled DNA stretch moves away from the fork, as exemplified by PCNA. In line with the pulse-only experiment, DEK was found to bind at later time points, with relative Western Blot signal intensities increasing to significance at 15 to 30 min after thymidine addition. Here too, DEK behaved similarly to histone H3. From these experiments we conclude that DEK is not a component of the active replisome but rather binds to DNA as it assembles into mature chromatin.

**Fig 3.**
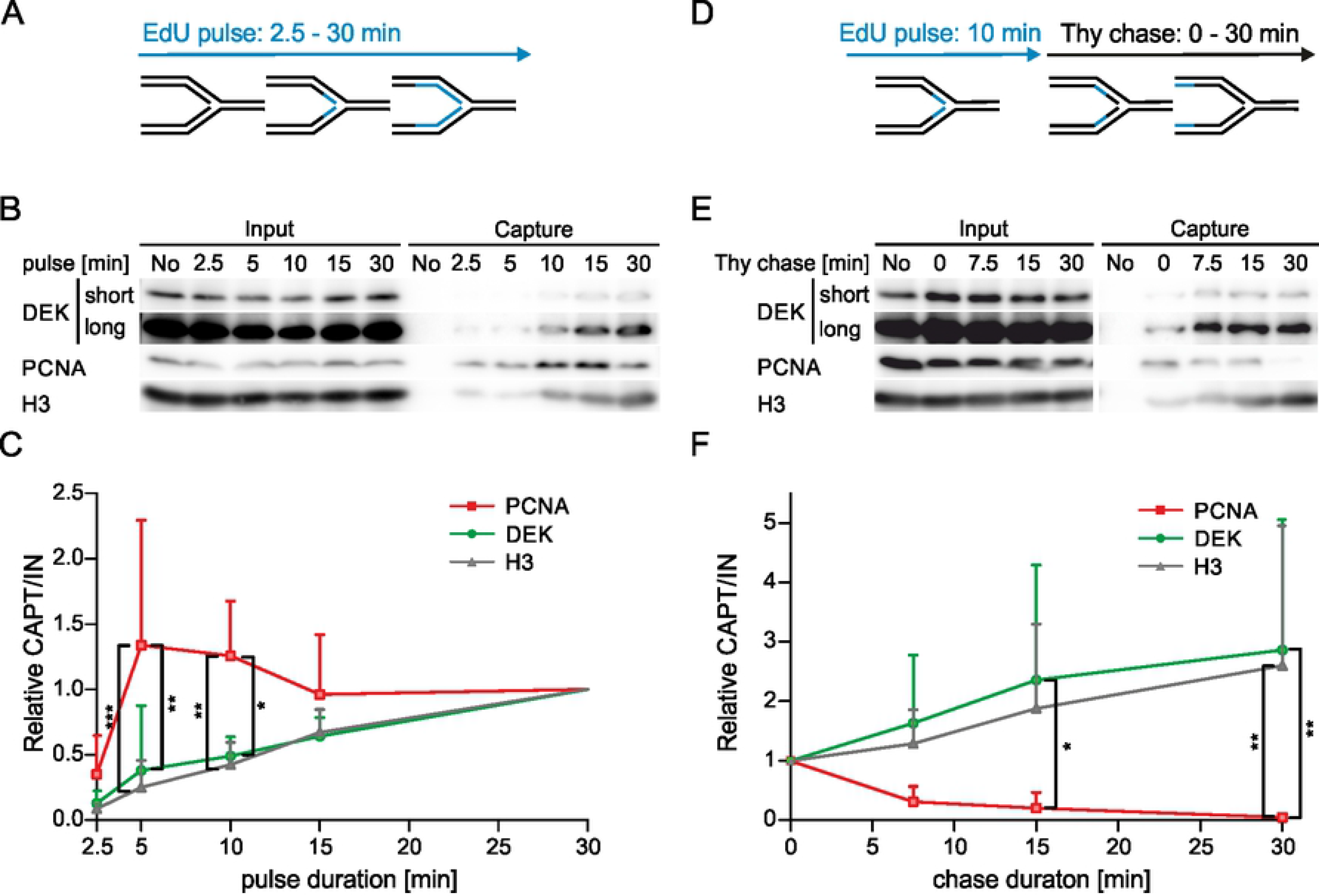
DEK is not a component of the replisome. (A-C) HeLa S3 cells were pulse-labelled with EdU for 2.5 – 30 min and biotin-azide was covalently attached via click chemistry. After cell lysis, EdU-biotin containing DNA fragments were precipitated using streptavidin-coupled magnetic beads. Bound proteins (capture) were identified by Western blot analysis. (A) Scheme of iPOND pulse experiment. (B) Representative Western blots of input and capture samples using antibodies specific for DEK, PCNA and histone H3. (C) Densitometric analysis. The fold change of captured protein is displayed relative to the value of the 30 min time point. Band intensities of capture samples were normalized to the respective input smaples. Shown are mean values from five independent experiments. One-sided error bars represent the S.D. 2way ANOVA with Bonferroni posttest: * p≤0.05, ** p≤0.01, *** p≤0.001. (D-F) HeLa S3 cells were pulse-labelled with EdU for 10 min, followed by a chase into thymidine containing medium for 0 – 30 min. iPOND was performed as in (A-C). (D) Scheme of iPOND chase experiment. (E) Representative Western blots of input and capture samples using antibodies specific for DEK, PCNA and histone H3. (F) Densitometric analysis. The fold change of captured protein is displayed relative to the value of the 0 min time point. Band intensities were normalized as in (C). Shown are mean values from five independent experiments, one-sided error bars represent the S.D. 2way ANOVA with Bonferroni posttest: * p≤0.05, ** p≤0.01, *** p≤0.001.

To complement this biochemical approach we studied the localization of DEK in replicating cells by superresolution microscopy (Fig 4). To this purpose, we took advantage of a U2-OS knock-in cell line expressing GFP-DEK from its endogenous promoter (Vogel et al, in preparation). Firstly, we employed structured illumination microscopy [52] and combined EdU labelling of nascent DNA with immunolabeling of PCNA. The images showed that DEK does not colocalize with sites of active replication (Fig 4 A). This finding was corroborated by stochastic optical reconstruction microscopy (STORM, Fig 4 B) [53]. This approach offers a key chance to investigate the distribution of nuclear proteins at the nanoscale level [54]. Also at this higher resolution, DEK is not found colocalizing with PCNA. Presently, we cannot exclude that the localization of DEK with respect to active replication foci may vary during S-phase progression, thus accounting for a partial enrichment of DEK in late chromatin fractions in iPOND experiments. Altogether, based on these data, we can exclude that DEK is part of the replisome making it very unlikely that its function in promoting replication fork progression occurs via a direct interaction.

**Fig 4.**
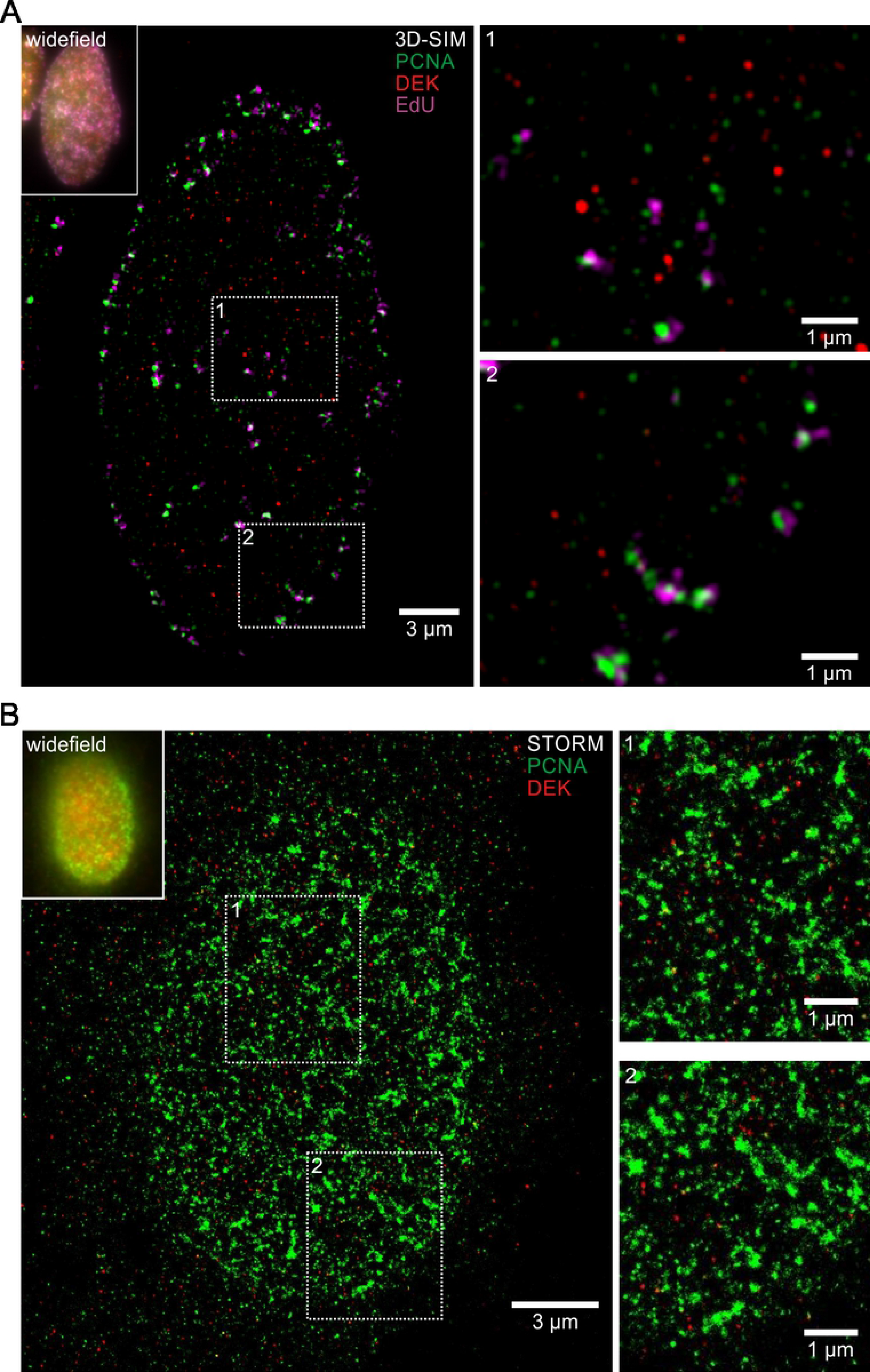
DEK does not colocalize with replication foci in superresolution images. (A) 3D-SIM superresolution microscopy images of DEK, EdU and PCNA distribution in early/mid S-phase. U2-OS GFP-DEK cells were treated with EdU for 10 min to label nascent DNA via click chemistry, and PCNA was visualized via indirect immunofluorescence. Shown is a single z-slice from the super-resolved image stack with two magnified insets. Red: GFP-DEK, green: PCNA (green), magenta: EdU. Upper left corner: Pseudo-widefield representation of the same nucleus by superimposition of all z-slices. (B) STORM superresolution microscopy images of DEK and PCNA distribution in early S-phase. DEK (red) and PCNA (green) were visualized via indirect immunofluorescence in U2-OS GFP-DEK cells with Alexa405/Alexa647 photoswitchable dye pairs respectively CF568. Shown is a single z-slice with two magnified insets. Top left corner: Widefield image of the same nucleus.

### Fork restart impairment observed under PARP1/2 inhibition depends on DEK expression

As we had observed that the effect of PARP1/2 inhibition on mildly impaired replication forks depended on DEK expression, we asked whether DEK levels would also impinge on the recovery of replication forks after stalling. PARP1/2 was shown to protect replication forks stalled by HU treatment and promote their effective restart [8, 9, 55], providing a suitable experimental paradigm to evaluate the effect of DEK downregulation.

We performed DNA fiber assays in shDEK and control cells in which forks were completely blocked using a prolonged exposure to a high dose of HU (4 mM for 4 h, see also S3 Fig), followed by removal of HU and release in fresh medium in the presence and absence of ABT-888 (Fig 5 A). Downregulation of DEK alone had no effect on the resumption of DNA synthesis after removal of HU, with about 80% of forks restarting in both control and shDEK cells. In the former, PARP1/2 inhibition led to a marked decrease of fork restart efficiency to about 60%, in line with published results [8]. In contrast, shDEK cells were completely protected from restart impairment, displaying a slightly higher number of restarting forks as compared to control cells not exposed to ABT-888 (Fig 5 C). These data reflect the same phenotype of DEK downregulation counteracting PARP1/2 inhibition as observed in the context of fork slowing by CPT, and underscore its functional relevance.

**Fig 5.**
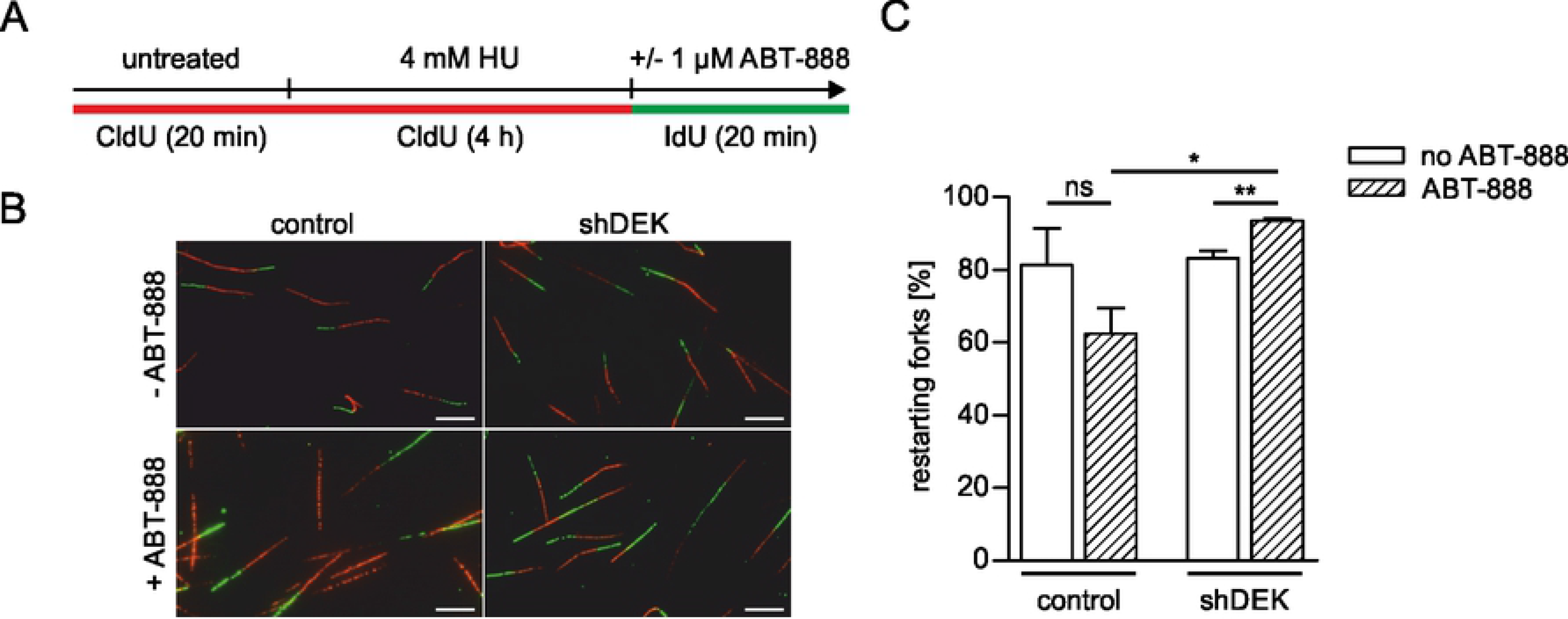
DEK depletion counteracts fork restart impairment due to PARP1/2 inhibition. (A-C) U2-OS control and shDEK cells were pulse-labelled with CldU for 20 min, followed by incubation with 4 mM HU for four hours to arrest replication forks. Forks were released in fresh IdU-containing medium in the presence or absence of 1 μM ABT-888. (A) Scheme of the fork restart experiment. (B) Representative confocal images for each experimental condition. CldU-labelled tracts were visualized in the red channel, IdU-labelled tracts in the green channel. Scale bar: 5 μm. (C) Quantification of results. The mean percentage of restarting forks from three independent experiments is shown. At least 300 fiber tracts were scored per experimental condition. Error bars represent the S.E.M. t-test: ns: not significant, * p≤0.05, ** p≤0.01. ABT-888 treated cells: hatched bars.

Fork impairment and stalling by HU treatment has been shown previously to elicit the robust formation of RPA-positive foci [56], reflecting RPA binding to single stranded DNA (ssDNA) which is extensively generated when polymerase and helicase activity are uncoupled. RPA protects this ssDNA from nucleolytic attack and serves multiple important functions in the repair and restart of damaged forks [57, 58]. To further investigate the effect of DEK expression on impaired replication forks we determined the formation of RPA-positive foci under conditions of replication stress combined with PARylation inhibition (Fig 6). We applied 2 mM HU for 80 min, as this dose and time of exposure was shown to elicit a maximal HU response in U2-OS cells [56], and measured RPA foci in EdU positive cells (see also S4 Fig for the quantification method). As observed for fork slowing and DSB formation, shDEK cells showed a significantly increased RPA response as compared to control cells. In the latter, additional PARP1/2 inhibition reduced the formation of RPA foci with respect to HU treatment only. This finding is in line with data from Bryant et al. [8] who reported reduced RPA foci in PARP-inhibited cells. Again, shDEK cells reacted differently, displaying a small, but significant increase in RPA-positive cells when exposed to HU in combination with ABT-888 (Fig 6 B). These data suggest that DEK plays a role in limiting the formation of long ssDNA stretches upon stalling or collapse of replication forks, and that its downregulation compensates for the previously described requirement for PARP1/2 activity for RPA binding at a subpopulation of stalled forks. These results further corroborate the existence of a reciprocal functional link between DEK and PARP1/2 in the response to replication stress.

**Fig 6.**
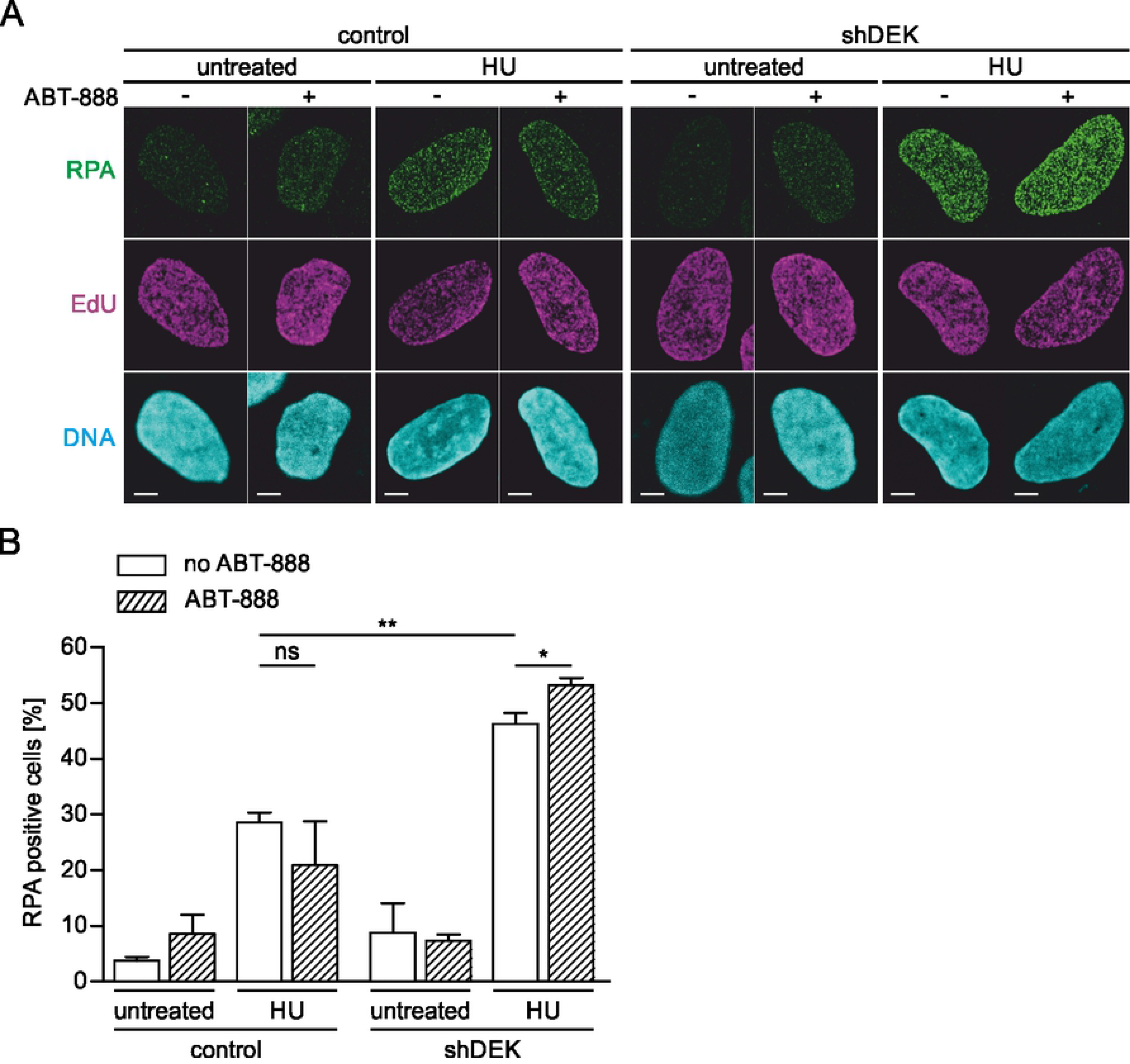
DEK counteracts the effect of ABT-888 on RPA foci formation under HU treatment. (A-B) U2-OS control and shDEK cells were pulse-labelled with EdU, then either left untreated or treated with 2 mM HU for 80 min in presence or absence of 1 μM ABT-888. RPA foci (green) were visualized via indirect immunofluorescence, EdU (magenta) using click chemistry. DNA was counterstained with Hoechst 33342 (cyan). (A) Representative confocal images for each experimental condition. Scale bar: 5 μm. (B) Percentage of RPA positive S-phase cells as determined using the automated foci counter of the BIC macro tool box (see also S4 Fig). Mean values from three independent experiments are shown. At least 97 cells were scored per experimental condition. Error bars represent the S.E.M. t-test: *p≤0.05, **p≤0.01.

### A DEK mutant with impaired PAR-interaction ability counteracts the effect of DEK downregulation on the response to replication fork stalling

In our previous work we described that DEK is modified by PAR covalently and non-covalently [18, 20, 49]. Interestingly, DEK shows a remarkably high affinity for long PAR chains exceeding that of histone H1. Therefore, we hypothesized that noncovalent DEK-PAR interaction would be important to mediate the effect of DEK on challenged replication forks. To verify this hypothesis, we sought to obtain a PAR-binding deficient mutant of DEK. We performed a systematic mutational study of the three previously described PAR-binding domains (PBDs) in the DEK primary sequence (PBD1: aa 158-181, PBD2: aa 195-222; PBD3: aa 329-352; Fig 7 A and S1 Table). Previous *in vitro* studies showed that they have different affinities for purified PAR. The strongest PAR binding domain in the DEK primary sequence is PBD2 at amino acid positions 195-222, partially overlapping with the SAP-box of DEK, which is its major DNA-binding domain [18]. Mutant peptides corresponding to the three PAR-binding domains of DEK were subjected to PAR-overlay assays to assess the effect of single and multiple amino acid exchanges on PAR binding *in vitro* (Fig 7 B, C; S5 Fig). We were able to identify mutations within the high-affinity PBD2 peptide which completely abrogated non-covalent PAR interaction (Fig 7 B, C, Mut1-3). Moreover, when peptides corresponding to the three PBDs were incubated simultaneously with purified PAR in the same slot blot, only peptide 195-222 gave rise to PAR-specific signals, suggesting that this domain is the predominant PAR-acceptor in DEK, outcompeting the weaker PBDs (data not shown). We then generated purified recombinant DEK carrying a mutated PBD2 (Mut2) and tested the effect of the identified mutations in the context of the full-length protein (Fig 7 D). By densitometric analysis we observed a reduction in PAR-binding affinity of about 50% as compared to the corresponding wildtype DEK sequence, confirming the data obtained with the isolated peptides. The overall DNA binding ability of the PBD2-Mut2 mutant was only slightly reduced as compared to the wildtype DEK. A band shift in EMSA assays became detectable at a molar ratio of DEK:DNA of 112 instead of 84. At higher molar ratios the binding behaviour of the two proteins was undistinguishable (Fig 7 E).

**Fig 7.**
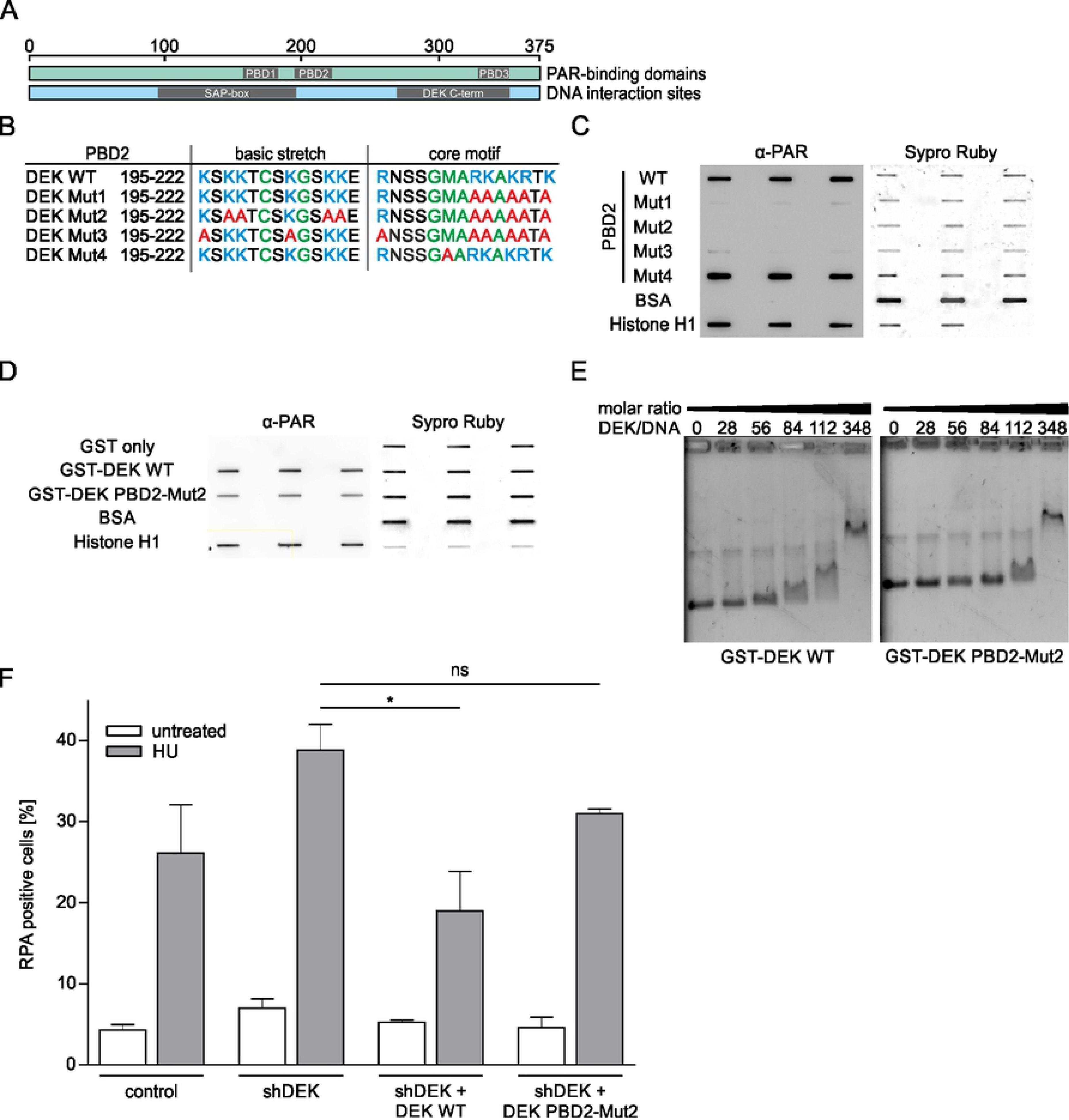
Non-covalent interaction of DEK with PAR is important for RPA foci formation upon HU treatment. (A) Schematic of the DEK protein with PAR-binding domains and DNA interaction sites. (B) Mutational analysis of PBD2 using recombinant peptides. (B) Basic (green) and/or hydrophobic (blue) amino acids were exchanged for alanine (red) as indicated. (C) Peptides were analysed in a PAR overlay assay to assess PAR-binding. PAR was detected by chemiluminescence using a specific antibody (α-PAR-10H). Equal membrane loading of peptides was verified using Sypro Ruby. One representative blot is shown. The experiment was performed in triplicate with similar results. (D) Recombinant full-length GST-DEK WT and GST-DEK PBD2-Mut2 were purified from *E.coli* and analysed using PAR-overlay assays. One representative blot out of two replicates is shown. (E) Analysis of the DNA binding ability of GST-DEK WT and GST-DEK PBD2-Mut2 via EMSA. Recombinant proteins were incubated with plasmid DNA in increasing molar ratios. DEK/DNA complexes were separated on agarose gels and visualized using ethidium bromide. One representative gel out of two replicates is shown. (F) Analysis of RPA foci formation. U2-OS shDEK cells were transfected with plasmids encoding GFP fused to either WT-DEK or DEK PBD2-Mut2. GFP-positive, low-level expressing cells were isolated by FACS. U2-OS control and shDEK cells expressing GFP-DEK fusion proteins were treated with 2 mM HU for 80 min. RPA foci formation was analysed by immunofluorescence as described in Fig 6, S-phase cells were identified by EdU labelling. The percentage of RPA positive S-phase cells was determined using the automated foci counter of the BIC macro tool box. The mean values from three independent experiments are shown. At least 100 cells were scored per experimental condition. Error bars represent the S.E.M. t-test: ns: not significant, * p≤0.05.

We then used the PDB2-Mut2 mutant to analyse the influence of PARylation of DEK on the formation of RPA foci upon fork stalling. Wildtype or PBD2-Mut2-DEK fused to GFP were expressed in shDEK and control cells and the number of RPA-positive cells after HU treatment was determined as above. Both GFP fusion proteins were expressed at comparable levels as verified by Western blot (S6 Fig).

Treatment with 2 mM HU robustly triggered RPA foci formation, to a higher level in shDEK cells as compared to controls, as already observed (Fig 7 F). Re-expression of WT DEK abrogated this effect confirming its specificity and reducing the number of RPA-positive cells to a level below that of HU-treated control cells. Importantly, the DEK mutant with reduced PAR-binding ability was much less effective in counteracting RPA-foci formation. We conclude from this result that the increase in HU-induced RPA foci mediated by DEK requires its non-covalent interaction with PAR. We cannot rule out, however, that other mechanisms and/or PAR-binding domains of DEK may be involved too, because the PBD2-Mut2 did not fully restore the level of RPA foci obtained in shDEK cells after HU treatment (Fig 7 F). Taken together, these data strongly suggest that the non-covalent interaction with PAR plays a major role in regulating how DEK affects the response to replication stress providing a first mechanistic insight in the complex molecular interplay of PARP1/2 and DEK.

## DISCUSSION

In this study we have explored a potential functional relationship between DEK and PARP1/2 in the context of DNA replication stress. For both proteins, there is consistent evidence for their involvement in the response to impaired DNA replication. Both DEK and PARP1/2 preferentially bind to unconventional non-B DNA structures like cruciform and G4 DNA [36, 59–61]. These structures are difficult to replicate and particularly abundant in heterochromatin. Both DEK and PARP1/2 are found enriched in chromatin of S-phase cells [62–65] and have been associated with the formation and maintenance of heterochromatin [39, 66]. DEK was shown to modulate the efficiency of DNA replication *in vitro* [42], and, more recently, we showed that normal DEK levels are necessary to sustain replication fork progression and to prevent fork rearrangements in cells undergoing replication stress [43]. DEK is a target for covalent modification by and non-covalent interaction with PAR. Covalent PARylation was reported to occur at glutamic acid 136 [67] and 207 [68], arginine 208 [68] and, most recently, at serine 279 [69]. Based on sequence alignment, DEK was further proposed to harbour three non-covalent PAR-binding domains [18], of which the central one (aa position 195-222) shows the strongest binding affinity and mediates about 50% of the PAR-binding activity of the protein *in vitro* (this study and [70]).

Impaired replication forks can activate PARP1/2 and PAR has been involved in the regulation of different types of fork processing and rearrangements. Thus, PARP1/2 can protect replication forks from extensive Mre11-dependent resection after HU treatment [8, 9], or from untimely resolution of RecQ-mediated reversal in cells treated with low doses of CPT [11]. Massive accumulation PAR, on the other hand, has adverse consequences. HU-induced prolonged fork stalling in cells with downregulated PAR glycohydrolase (PARG) leads to fork collapse and DSB formation [71]. The molecular events orchestrated by PARP1/2 activation during DNA replication therefore seem to depend, in a yet poorly understood fashion, on the type and extent of the replication problem, which in turn determine the amount and possibly also the structure of the polymer formed. Consequently, inhibiting PARylation during DNA replication may have different outcomes, depending not only on the dose and the duration of the inhibitor treatment but also on the status of the replication machinery. As shown here, short term inhibition of PARP1/2 using ABT-888 aggravates fork retardation and DNA damage induced by mild replication stress, while long term exposure to AZD-2281 was shown to accelerate fork speed both in the presence and in the absence of DNA replication inhibitors [10, 72]. Our data on fork progression in shDEK cells suggest that challenged replication forks can switch between two opposing responses to PARP1/2 inhibition depending on the level of DEK expression. Interestingly, the restoration of fork speed observed in CPT-treated, PARP-inhibited shDEK cells was accompanied by a reduction in the level of replication-associated DSBs. This is in agreement with the finding that fork acceleration activates the DNA damage response only if fork speed exceeds a critical threshold [72], and poses an argument against replication fork runoff occurring in CPT-treated shDEK cells upon treatment with ABT-888. The behaviour of forks stalled by high doses of HU followed a similar pattern with respect to PARP-inhibition and DEK downregulation as CPT-induced fork slowing, being most efficiently restored in PARP-inhibited shDEK cells, which is suggestive of a common mechanism.

How DEK affects the sensitivity of replication forks towards PARP inhibition, is a matter of speculation so far. Based on our iPOND and superresolution microscopy data it seems unlikely that DEK acts directly at the fork, since we did not find it associated to nascent DNA before the stage of nucleosome formation. Rather, the data presented here lend credit to the hypothesis that DEK is part of a DNA replication regulatory circuitry orchestrated by PAR that affects response to acute PARPi treatment. Recently, a fork speed regulatory network has been proposed which controls replication fork progression via PARylation and the p53-p21 axis [72]. Both p21 and PAR act as suppressors of fork speed in an interdependent manner, as PARP1 additionally represses p21 expression. Intriguingly, downregulation of DEK expression was shown to result in p53 stabilization and increased p21 levels [32], alterations which may indirectly affect the response of fork speed to PARP1/2 inhibitors. Since a PAR-binding defective mutant of DEK partially rescued fork restart impairment by PARP inhibition we cannot exclude that DEK exerts its influence also by directly participating in the PARylation mediated sensing of replication problems. Altogether, this study pinpoints DEK as an important mediator of the PARP1/2-dependent response of replicating cells to fork impairment, a previously unrecognized function of DEK which has implications for tumor therapy and warrants further investigation.

## Supporting information

Supporting Information Ganz_Vogel et al

## ACKNOWLEDGEMENTS

We thank Eva Gwosch for providing U2-OS GFP-DEK cells, and Aswin Mangerich, Martin Stöckl and the whole BIC team for fruitful discussions. Further, we thank Anja Deutzmann for initial input in this study, and the FlowKon core facility at the University of Konstanz for cell sorting.

## SUPPORTING INFORMATION

**S1 Table Primer sequences for the site-directed mutagenesis of the DEK primary sequence**

**S1 Fig. AZD-2281 but not ABT-888 induces DNA damage in U2-OS cells**

**S2 Fig. Absence of combined positive effect of DEK downregulation and PARP1/2 inhibition on fork progression under high doses of HU**

**S3 Fig. HU-induced fork arrest**

**S4 Fig. Determination of RPA-positive cells**

**S5 Fig. Mutational analysis of PBD1 and PBD3 using recombinant peptides S6 Fig. Reconstitution of U2-OS shDEK cells with DEK WT-GFP or DEK PBD2-Mut2-GFP**

