## Supporting Information Ganz_Vogel et al for "The oncoprotein DEK affects the outcome of PARP1/2 inhibition during replication stress"

S1 Table Primer sequences for the site-directed mutagenesis of the DEK primary sequence

| | |
|---|---|
| PBD2-Part1-fwd | GCCGAAATCTGCAGCAACTTGTAGCAAAGGCAGTAAAAAGGAACGG |
| PBD2-Part1-rev | CTGCCTTTGCTACAAGTTGCTGCAGATTTCGGCAATGGTTTGC |
| PBD2-Part2-fwd | GCAAAGGCAGTGCAGAACGGAACAGTTCTGGAATGGCAAGG |
| PBD2-Part2-rev | GTTCCGTTCTGCTGCACTGCCTTTGCTACAAGTTGCTGCAG |
| PBD2-Part3-fwd | CTGGAATGGCAGCGGCGGCTAAGCGAACCAAATGTCCTG |
| PBD2-Part3-rev | GGTTCGCTTAGCCGCCGCTGCCATTCCAGAACTGTTCCG |
| PBD2-Part4-fwd | GGAATGGCAGCGGCGGCTGCGGCAACCGCATGTCTTCTTGAAATTCTG |
| PBD2-Part4-rev | GGACATGCGGTTGCCGCAGCCGCCGCTGCCATTCCAGAAC |
| shDEK-Part1-fwd | CCAGTGCGAACCTCGAGGAAGTCACAATGAAACAGATTTGC |
| shDEK-Part1-rev | CATTGTGACTTCCTCGAGGTTCGCACTGGCCAGTAATTTC |
| shDEK-Part2-fwd | CTGGCAAGCGCGAATCTTGAGGAGGTCACAATGAAACAG |
| shDEK-Part2-rev | GTGACCTCCTCAAGATTCGCGCTTGCCAGTAATTTCTTTATTG |

**S1 Fig.**

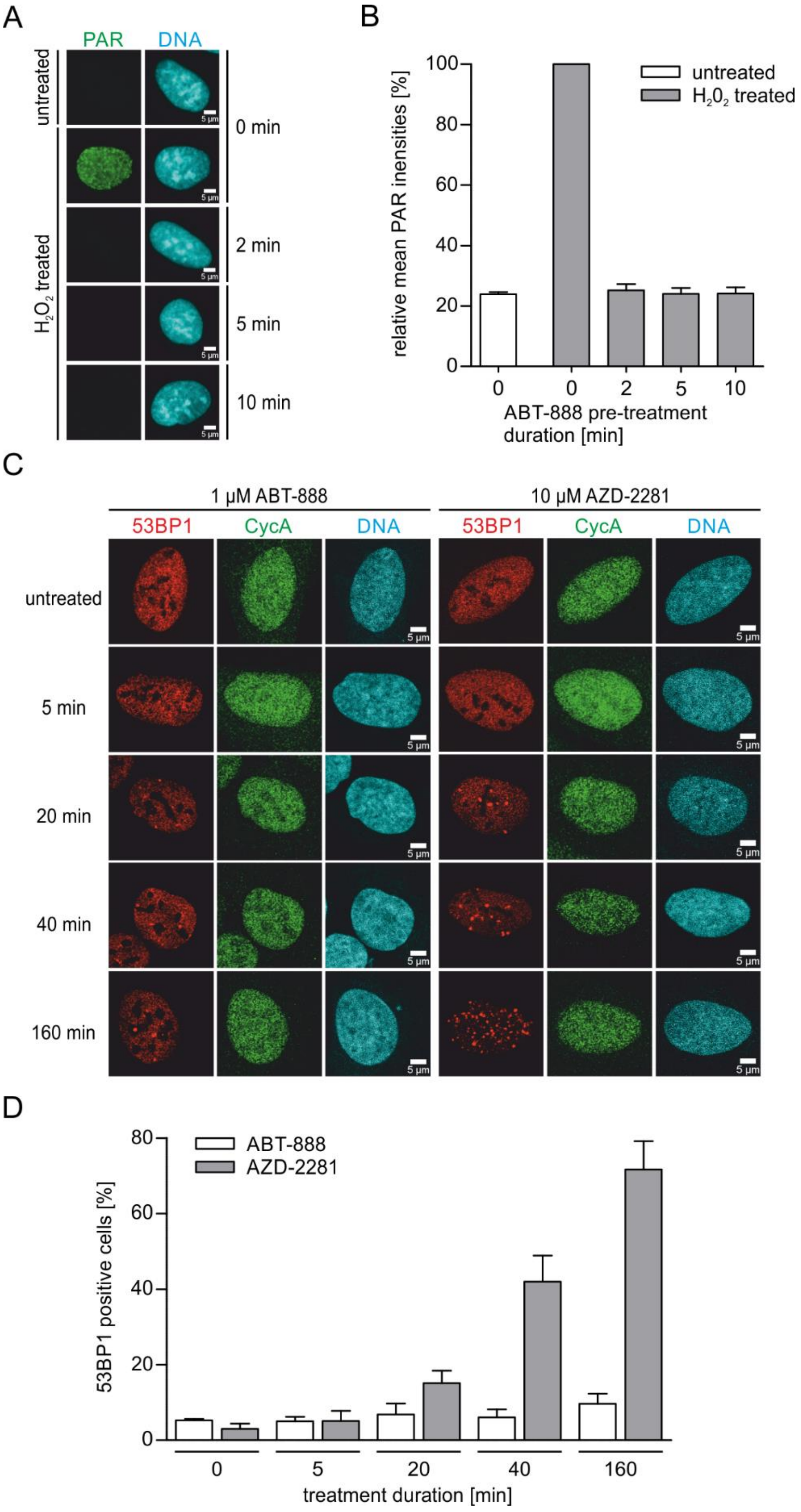
A
PAR
DNA
untreated
H2O2 treated
0 min
2 min
5 min
10 min
5 µm
B
relative mean PAR inensities [%]
untreated
H202 treated
ABT-888 pre-treatment duration [min]
0
0
2
5
10
C
1 µM ABT-888
10 µM AZD-2281
53BP1
CycA
DNA
untreated
5 min
20 min
40 min
160 min
D
53BP1 positive cells [%]
ABT-888
AZD-2281
treatment duration [min]
0
5
20
40
160

**S1 Fig. AZD-2281 but not ABT-888 induces DNA damage in U2-OS cells**

(A-B) Inhibition of PARP1/2 activity by ABT-888. U2-OS cells were pre-treated with 1 µM ABT-888 for increasing periods of time or left untreated. PAR formation was induced with 800 µM $H_2O_2$ for 10 minutes and visualized via indirect immunofluorescence using a specific antibody (PAR-10H, green). DNA (cyan) was counterstained using Hoechst 33342. (A) Representative microscopic images are shown. (B) Quantification of PAR fluorescence intensities. Values were normalized to t=0 min after $H_2O_2$ treatment in presence of the inhibitor. At least 100 nuclei were imaged per experimental condition. The experiment was performed in triplicates. Mean values are shown. Error bars show the S.E.M. Untreated: white bar, $H_2O_2$-treated: grey bars. (C-D) Induction of DNA strand breaks in the presence of ABT-888 and AZD-2281. U2-OS cells were either treated with 1 µM ABT-888 or 10 µM AZD-2281 for increasing periods of time or left untreated. 53BP1 foci formation (red) and cyclin A expression (green) were visualized via indirect immunofluorescence. DNA (cyan) was counterstained with Hoechst 33342. (C) Representative confocal immunofluorescence images for each experimental condition are shown. (D) Quantification of results. The mean percentage of 53BP1 positive S-phase cells is shown. The amount of foci was quantified using the foci counter tool of the BIC macro tool box. At least 300 nuclei were imaged per experimental condition. The experiment was performed in triplicates. Mean values are shown. Error bars show the S.E.M. ABT-888-treated: white bars, AZD-2281-treated: grey bars.

**S2 Fig.**

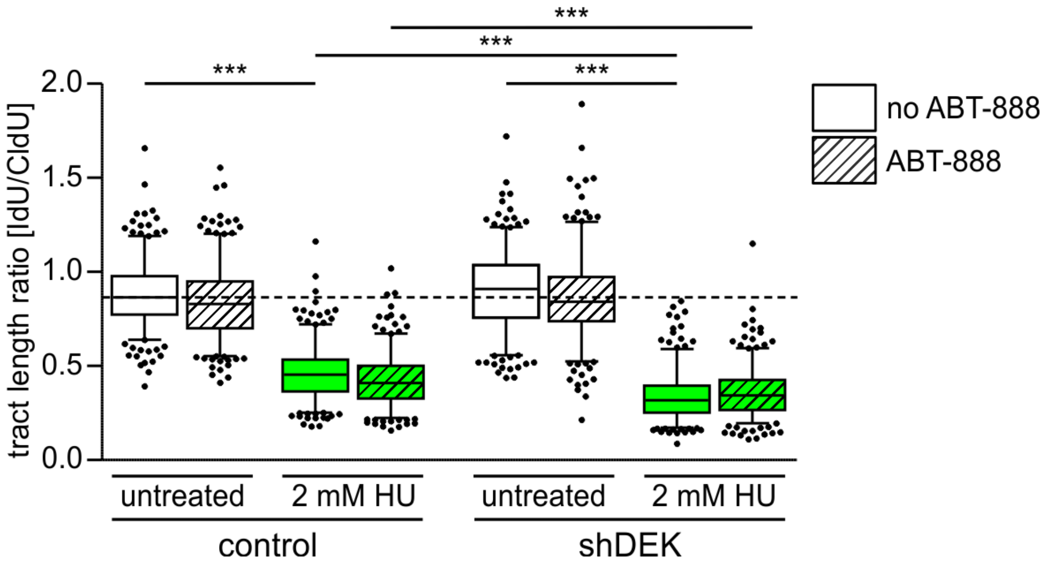


**S2 Fig. Absence of combined positive effect of DEK downregulation and PARP1/2 inhibition on fork progression under high doses of HU.**

A DNA fiber assay was performed as shown in Fig 1. U2-OS control and shDEK cells were pulse-labelled with CldU for 20 min, followed by incubation with IdU for 20 minutes in the presence or absence of 2 mM HU and 1 μM ABT-888. Thymidine analogues were visualized via indirect immunofluorescence after fiber spreading. A total of 300 tracts for each experimental condition were scored. The experiment was performed in triplicates. The bands inside the boxes display the median, whiskers indicate the 5th to 95th percentile and black dots mark outliers. t-test: *** p≤0.001. ABT-888 treated cells: hatched bars.

**S3 Fig.**

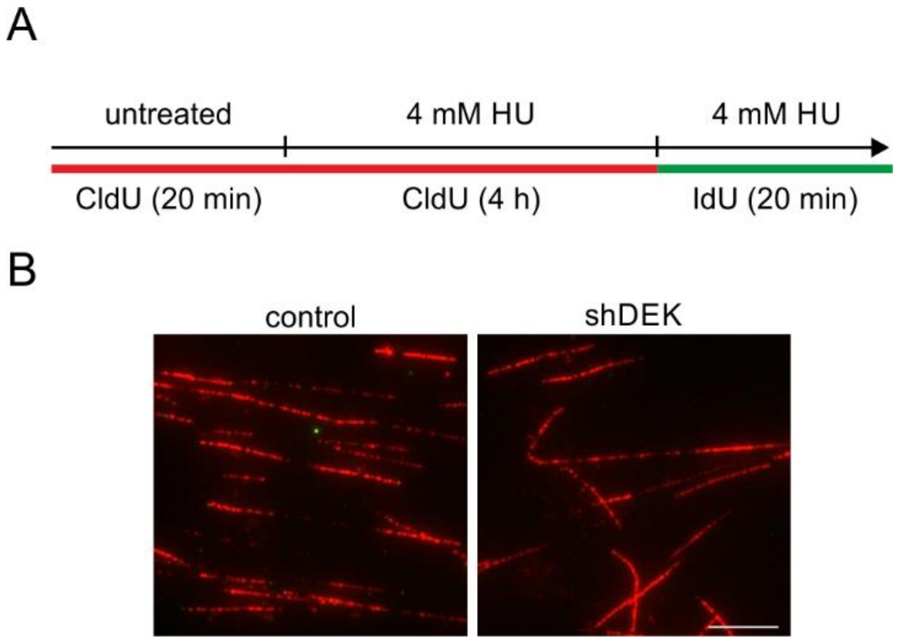


**S3 Fig. HU-induced fork arrest**

(A) Scheme of the control experiment for the fork restart assay shown in Fig 5. U2-OS control and shDEK cells were pulse-labelled with CldU (red) for 20 min, followed by incubation with 4 mM HU for four hours to arrest replication forks. Fork arrest was maintained in fresh IdU-containing (green) medium in the presence of 4 mM HU. (B) Representative images of DNA fiber spreads from U2-OS control and shDEK cells treated as in A. Scale bar: 5 µm.

**S4 Fig.**

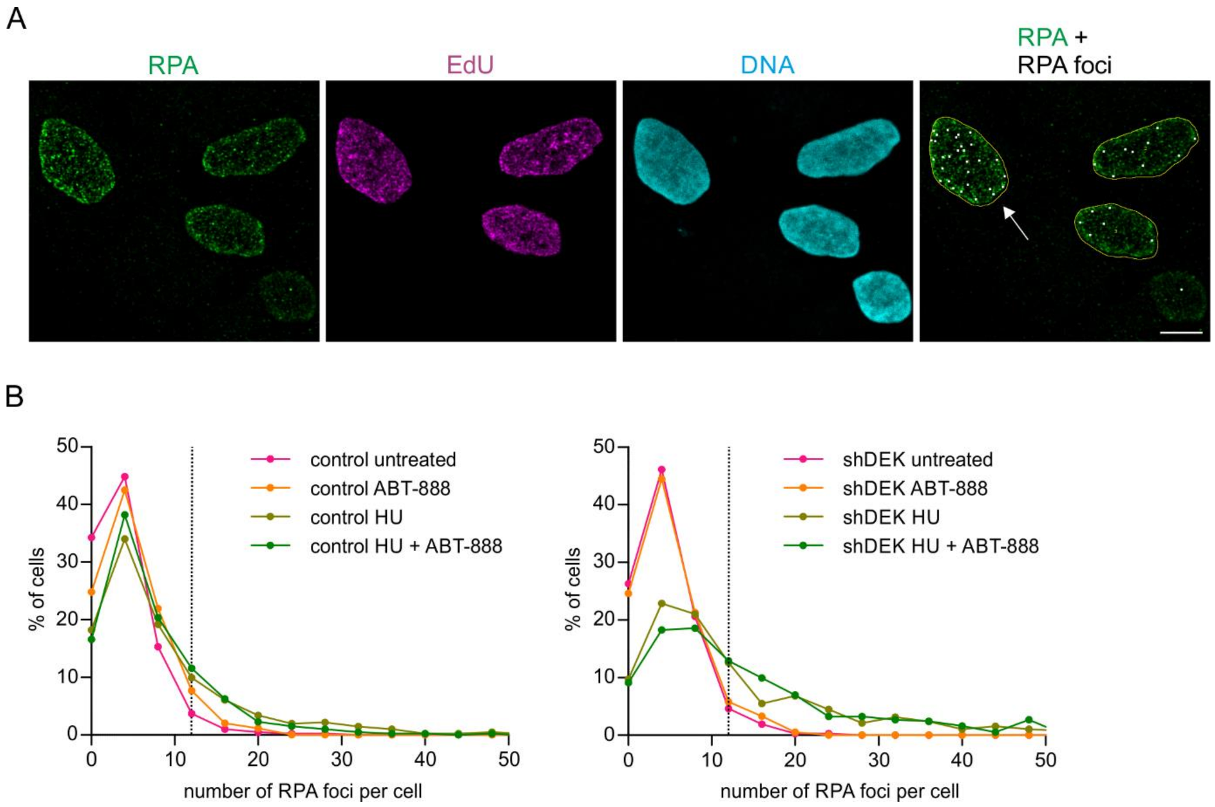


### S4 Fig. Determination of RPA-positive cells

(A) Representative confocal image showing HU-induced RPA foci in nuclei of U2-OS cells. The experiment was performed as described in Fig 6. RPA foci were detected using the automated foci counter of the BIC macro tool box. The right image shows detected foci (white dots) superimposed on the RPA fluorescence signal. Only the left cell (white arrow) exceeded the threshold of 11 foci set for this experiment and was classified as RPA positive. Scale bar: 10 µm. (B) Histograms of RPA foci distribution in S-phase cells. Cells were treated as described in Fig 6 A. Left panel: U2-OS control cells, right panel: U2-OS shDEK cells. The dashed line marks the threshold for RPA positive cells. Bin width: 4 RPA foci per cell.

**S5 Fig.**

A

| PBD1 | | basic stretch | core motif |
|---|---|---|---|
| DEK WT | 158-181 | KSICEVLDLERS | GVNSELVKRILN |
| DEK Mut1 | 158-181 | KSICEVLDLERS | GVNSELVAAILN |
| DEK Mut2 | 158-181 | KSICEVLDLERS | GANSEAAKRAAN |
| **PBD3** | | **basic stretch** | **core motif** |
| DEK WT | 329-352 | IKKLLASANLEE | VTMKQICKKVYE |
| DEK Mut1 | 329-352 | IKKLLASANLEE | VTMAQICAAVYE |
| DEK Mut2 | 329-352 | IKKLLASANLEE | VTMKQAAKKAAE |

B

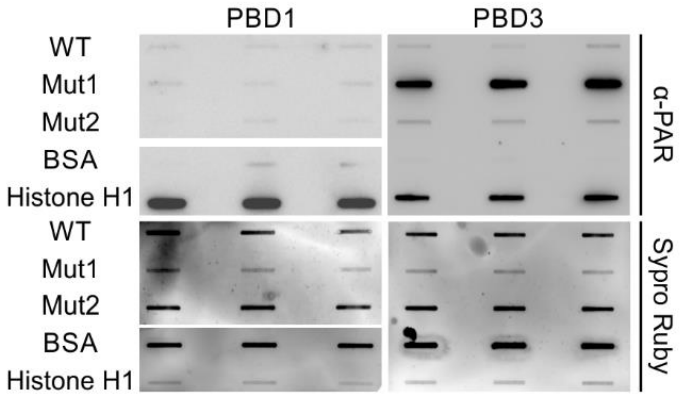


### S5 Fig. Mutational analysis of PBD1 and PBD3 using recombinant peptides

(A) Basic (green) and/or hydrophobic (blue) amino acids were exchanged for alanine (red) as indicated. (B) Peptides were analysed in a PAR overlay assay to assess PAR-binding. PAR was detected by chemiluminescence using a specific antibody (PAR-10H). Equal membrane loading of peptides was verified using Sypro Ruby. One representative blot is shown. The experiment was performed in triplicate with similar results.

**S6 Fig.**

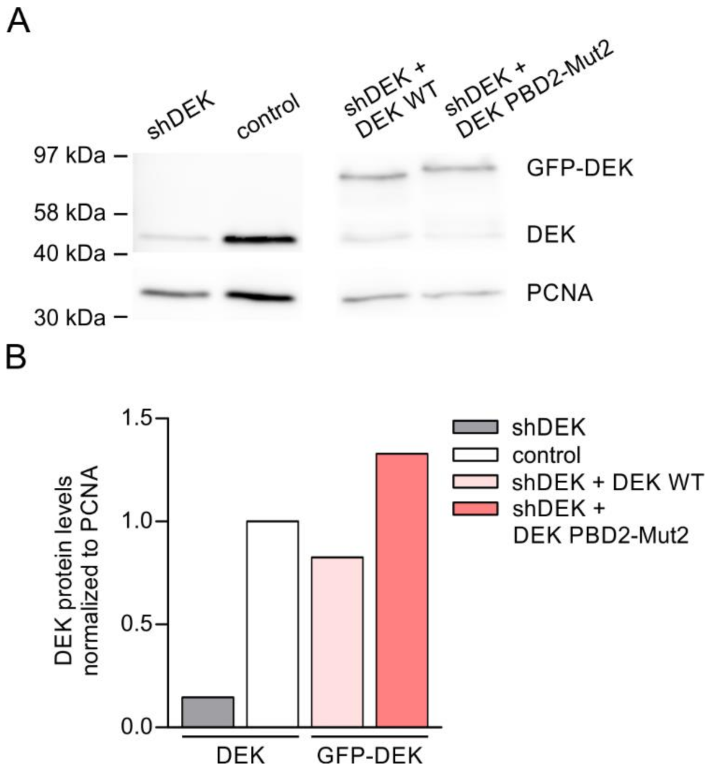


**S6 Fig. Reconstitution of U2-OS shDEK cells with DEK WT-GFP or DEK PBD2-Mut2-GFP**

U2-OS shDEK cells were transfected with plasmids encoding DEK WT-GFP or DEK PBD2-Mut2-GFP. GFP-positive, low level expressing cells were isolated using FACS. (A) Expression levels of ectopic DEK variants as well as endogenous DEK in U2-OS control and shDEK cells were visualized by Western blot. PCNA served as loading control. (B) Densitometric analysis. Corresponding DEK band intensities were normalized to PCNA and are displayed relative to endogenous DEK levels in control cells.
